# Inference of phenotype-defining functional modules of protein families for microbial plant biomass degraders

**DOI:** 10.1101/005355

**Authors:** S. G. A. Konietzny, P. B. Pope, A. Weimann, A. C. McHardy

**Affiliations:** Max-Planck Research Group for Computational Genomics and Epidemiology, Max-Planck Institute for Informatics, University Campus E1 4, 66123 Saarbrücken, Germany; Department of Chemistry, Biotechnology and Food Science, Norwegian University of Life Sciences, Post Office Box 5003, 1432 Ås, Norway; Department of Algorithmic Bioinformatics, Heinrich Heine University Düsseldorf, 40225 Düsseldorf, Germany

**Keywords:** Latent Dirichlet Allocation, LDA, probabilistic topic models, (ligno)cellulose degradation, plant biomass degradation, phenotype-based identification of functional modules, pectin degradation, probabilistic topic models, feature ranking, polysaccharide utilization loci, gene clusters

## Abstract

**Background:** Efficient industrial processes for converting plant lignocellulosic materials into biofuels are a key challenge in global efforts to use alternative energy sources to fossil fuels. Novel cellulolytic enzymes have been discovered from microbial genomes and metagenomes of microbial communities. However, the identification of relevant genes without known homologs, and elucidation of the lignocellulolytic pathways and protein complexes for different microorganisms remain a challenge.

**Results:** We describe a new computational method for the targeted discovery of functional modules of plant biomass-degrading protein families based on their co-occurrence patterns across genomes and metagenome datasets, and the strength of association of these modules with the genomes of known degraders. From more than 6.4 million family annotations for 2884 microbial genomes and 332 taxonomic bins from 18 metagenomes, we identified five functional modules that are distinctive for plant biomass degraders, which we call plant biomass degradation modules (PDMs). These modules incorporated protein families involved in the degradation of cellulose, hemicelluloses and pectins, structural components of the cellulosome and additional families with potential functions in plant biomass degradation. The PDMs could be linked to 81 gene clusters in genomes of known lignocellulose degraders, including previously described clusters of lignocellulolytic genes. On average, 70% of the families of each PDM mapped to gene clusters in known degraders, which served as an additional confirmation of their functional relationships. The presence of a PDM in a genome or taxonomic metagenome bin allowed us to predict an organism’s ability for plant biomass degradation accurately. For 15 draft genomes of a cow rumen metagenome, we validated by cross-linking with confirmed cellulolytic enzymes that the PDMs identified plant biomass degraders within a complex microbial community.

**Conclusions:** Functional modules of protein families that realize different aspects of plant cell wall degradation can be inferred from co-occurrence patterns across (meta)genomes with a probabilistic topic model. The PDMs represent a new resource of protein families and candidate genes implicated in microbial plant biomass degradation. They can be used to predict the ability to degrade plant biomass for a genome or taxonomic bin. The method would also be suitable for characterizing other microbial phenotypes.

## Background

Lignocellulose is an integral part of plant cell walls and is responsible for the structural integrity and robustness of crops, grasses and trees. The high energy content and renewability of lignocellulosic plant material make it a promising alternative energy resource, particularly for the production of biofuels [1, 2]. Industrial methods of degrading recalcitrant plant cell wall material remain inefficient [3], which has created great interest in lignocellulolytic microbial communities and their member organisms [4]. The genes of lignocellulolytic organisms represent a promising source of potential enzymes for improving the industrial degradation processes [4, 5]. Plant cell walls consist of cellulose and hemicelluloses (e.g. xylan, xyloglucan, β-glucan), which are cross-linked by lignin, and pectins [6, 7]. Cellulose is a macromolecule of β-(1,4)-linked D-glucose molecules. Xylans and β-glucans are homopolysaccharides composed of either xylose or β-1,3, β-1,4-linked D-glucose, respectively, and are commonly found in plant cell walls of grasses. Xyloglucan is a hemicellulose occurring in the plant cell wall of flowering plants and consists of a glucose homopolysaccharide backbone with xylose side chains, which are occasionally linked to galactose and fucose residues. Pectin is a heteropolysaccharide that represents a major component of the middle lamella of plant cell walls. Finally, lignin is a strongly cross-linked polymer of different aromatic compounds.

The degradation of lignocellulosic plant material requires the concerted action of different carbohydrate-binding modules (CBMs) and catalytic enzymes, such as cellulases, xylanases, pectin lyases and peroxidases [8–10]. The CAZy database [11] distinguishes four important subclasses of carbohydrate-active enzymes (‘CAZymes’); glycoside hydrolases (GHs), glycosyltransferases (GTs), polysaccharide lyases (PLs) and carbohydrate esterases (CEs). Carbohydrate-active proteins often have a modular composition, i.e. they possess a multi-domain architecture. Several multifunctional enzymes that combine different catalytic domains as well as one or more CBMs are found to be active in lignocellulose degradation [12]. Microorganisms use different strategies to degrade recalcitrant plant material. The ‘free enzyme’ and cellulosome strategies are the most widely used by known microbial plant biomass degraders [12, 13]. The ‘free enzyme’ paradigm is frequently employed by aerobic bacteria and involves the secretion of cellulolytic enzymes to degrade lignocellulose in the external medium. The cellulosome-based strategy has so far only been described for anaerobic bacteria [13]. Cellulosomes are large protein complexes that incorporate cellulolytic enzymes as well as CBMs for localized lignocellulose degradation [14]. The cellulosome includes a scaffoldin backbone to which cellulases and hemicellulases attach via cohesin–dockerin interactions. The corresponding (hemi)cellulases contain the dockerin domains, one or more catalytic domains (e.g. glycoside hydrolase enzymes) and non-catalytic CBMs [14]. More recently, two additional strategies for (hemi)cellulose degradation have been outlined. Sus-like protein systems rely on mechanisms that are similar to the starch utilization (Sus) system in *Bacteroides thetaiotaomicron* [15, 16], which are mediated by enzymes located in the outer membrane [17]. The second one involves the oxidative cleavage of cellulose by copper mono-oxygenases, a mechanism that increases the efficiency of the hydrolytic enzymes [18]. However, certain cellulolytic organisms, such as *Fibrobacter succinogenes* and *Cytophaga hutchinsonii*, do not seem to use any of the known mechanisms [13]. Additional insights into microbial degradation processes have been generated in studies of microbial communities using metagenomics. This has led to the identification of thousands of putative carbohydrate-active genes [19, 20] and several novel genes encoding proteins with cellulolytic activities from uncultured organisms [21–23]. Overall, more than 1000 cellulase genes have been discovered by genomic and functional screens [24]; however, important details about their microbial degradation mechanisms still remain unresolved [13, 25]. Therefore, the discovery of novel protein families that are involved in plant biomass degradation is still an ongoing effort.

The CAZYmes Analysis Toolkit (CAT) can be used to assign protein families from the CAZy database to protein sequences that have been annotated with Pfam, based on a set of pairwise association rules between CAZy and Pfam families [26]. CAT deduces its rules from the frequencies of modular proteins with Pfam and CAZy assignments in the CAZy database. However, this implies limitations for its application to families that have not yet occurred in the CAZy database. An alternative approach for determining protein families that participate in a particular process but have no homologs with known activities is to use genomic information in combination with other information sources. A functional context can be assigned to a new family by determining its association with a phenotype across a set of genomes. Depending on the granularity of the assigned context, this approach allows to narrow down the set of possible functions for an uncharacterized protein family. Applied to thousands of families on a large scale, this allows the *de novo* discovery of phenotype-defining protein families, genes or entire functional modules [27]. Several methods for ranking genes or pathways by their assumed relevance for a certain phenotype have been described [28–36]. These methods measure the association of individual protein families, known pathways or single nucleotide polymorphisms [34] with the presence or absence of phenotypes across a set of genomes. In some examples, the search space is limited to proteins in predicted operon structures [36] or pairs of functionally coupled proteins [35]. We have previously described a family-centric method for the identification of protein families involved in lignocellulose degradation [28]. This method uses an ensemble of linear L1-regularized Support Vector Machine classifiers trained with the genome annotations of known lignocellulose-and non-lignocellulose-degrading species. Other methods apply similar ranking approaches which are followed by a clustering step, where phenotype-associated families are grouped into modules based on their co-occurrence patterns across organisms, which are likely to indicate functional dependencies [29, 30]. However, we suggest that the order of steps should be reversed, because methods based on individual families have limitations: Proteins may have multiple functions (which is called ‘moonlighting’ [37]) and some families perform different functions within multiple functional modules (‘the patchwork hypothesis’ [38]). Consequently, proteins that are involved in lignocellulose degradation might also be active in other processes in non-lignocellulose-degrading organisms, which would reduce the global correlation pattern of their absence/presence profiles with the organisms’ ability to degrade lignocellulose. Moonlighting proteins might thus be missed. In contrast to approaches that focus primarily on single proteins, methods that target functional modules, i.e. sets of functionally related proteins that are jointly involved in a biological process such as lignocellulose degradation, have better chances of identifying moonlighting proteins, due to the functional dependencies between the families involved in the process.

Pathway-centric methods search for sets of functionally coupled protein families related to a specific phenotype. Often they use prior information about pathways from, for example, the KEGG [31] or BioPath [32] databases in the form of organism-specific enzyme reaction networks based on enzyme classification (EC) numbers. NIBBS (Network Instance-Based Biased Subgraph Search) searches for phenotype-associated edges in order to identify phenotype-related enzyme reactions in a KEGG-based network [33]. Similarly, MetaPath identifies subgraphs of a KEGG-derived network by assessing the statistical support of phenotype associations for every edge [31]. To date, there has been no application of pathway-centric methods to the study of lignocellulose degradation. Moreover, because of their focus on well-defined reaction networks, these methods have limitations for the analysis of metagenome samples, which often allow only partial metabolic reconstructions. Furthermore, species from newly sequenced microbial communities are likely to have a distinct metabolism from well-studied model species, and the latter have been the basis for most of the currently described reaction networks. We are not aware of a method for inferring phenotype-associated functional modules that is applicable to metagenomes and does not require prior knowledge about the underlying enzyme reaction networks or the target pathways. However, such a method would represent an important addition to computational metagenome analysis methods [39].

An indication of the functional context for a protein family can be obtained by clustering families by their co-occurrences across genomes [40, 41]. We have previously used Latent Dirichlet Allocation (LDA) [42], a Bayesian method, to infer 200 functional modules of protein families from 575 prokaryotic genomes [43]. The modules cover a broad range of biochemical activities, including several known protein complexes, metabolic pathways and parts of signal transduction processes. They show significant functional coherence according to high-confidence protein–protein interactions from the STRING database [44].

Here, we describe a method for determining the functional modules associated with microbial plant biomass degradation that uses LDA and, subsequently, selection of the relevant functional modules by the strength of their associations with the plant biomass degradation phenotype. The modules were learned from a large dataset of nearly 3000 sequenced bacterial and archaeal genomes and taxonomic bins of 18 metagenomes. Based on abundance estimates reported by Medie *et al*. [45] and Berlemont *et al*. [46], the relative abundance of species possessing plant biomass degradation capabilities within the sequenced genomes could exceed 20–25%; however, only a small set of species have been confirmed to date to possess such capabilities [4]. With our method, genomes of both known and unknown degraders could be included in the inference process and used to identify distinct sets of protein families that are specific for microbial plant biomass degraders. The use of metagenome data allows us to incorporate information from environmental communities into the inference process.

We identified five functional modules for plant biomass degradation, which we call PDMs. The PDMs included many protein families that are known to be involved in plant biomass degradation, and a substantial number of families that have not previously been linked to microbial plant biomass degradation. To verify the relevance of these newly identified PDMs and candidate families, we searched for gene clusters including the families of the PDMs. Several of the identified clusters are known to be active in the degradation of lignocellulose. Furthermore, the PDMs had a predictive value for identifying plant biomass degraders from the genomes of sequenced isolates or of plant biomass-degrading microbial communities.

## Results and discussion

We generated ∼6.4 million protein annotations with Pfam and CAZy families for 2884 bacterial and archaeal genomes from the Integrated Microbial Genomes database (IMG) and 332 taxonomic bins from 18 metagenomes (Methods). We then used a two-step approach to identify functional modules that are distinctive for microbial lignocellulose degraders: First, the annotated dataset was processed with LDA and 400 potential functional modules were inferred, where each corresponded to a set of Pfam and/or CAZy families (Figure 1, Steps 1 and 2). The modules were learned in an unsupervised fashion without consideration of the organisms’ phenotypes, as in [43]. In the second step, we ranked the 400 functional modules according to their strength of association with the genomes of plant biomass degraders across a subset of the genomes consisting of 38 known lignocellulose degraders and 82 non-degraders (Figure 1, Step 3). For this, we defined ‘genome-specific’ module weights, which corresponded to the fraction of a module’s protein families that were annotated for a certain genome or taxonomic bin (‘completeness scores’). Functional modules were considered to be present in a genome or bin if their completeness score reached a certain threshold. For each module, we determined the best setting for this threshold, corresponding to the one which optimally separated the genomes of degraders and non-degraders according to the F-measure (the weighted harmonic mean of precision and recall, Methods). The modules with the largest F-scores were strongly associated with the genomes of lignocellulose degraders, as indicated by an average F-score of 87.45% for the top ten modules.

**Figure 1:**
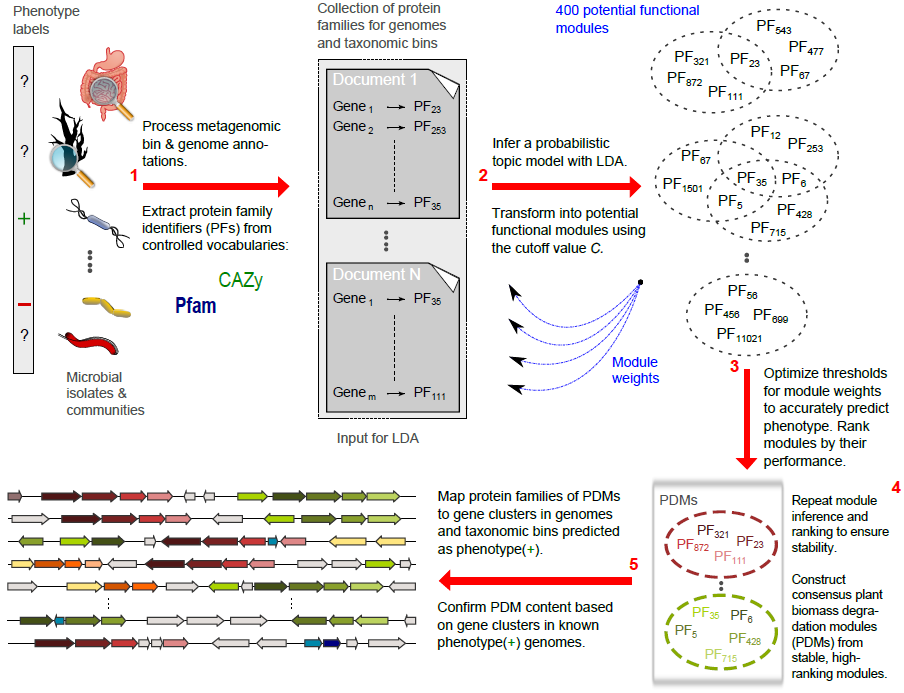
Identifying phenotype-related functional modules. We used protein sequences from 2884 prokaryotic isolate species and 18 microbial communities, some of which are known to be active in lignocellulose degradation. Known lignocellulose degradation abilities are indicated by phenotype labels (+/-). For the metagenomes, we only considered protein coding sequences with predicted taxonomic origins assigned by a taxonomic binning method (*PhyloPythia* or *PhyloPythiaS)*. We used HMMER to assign protein family annotations from Pfam and CAZy to all input sequences, and summarized the set of (meta-)genome annotations as a document collection for LDA (**1**). Each document is composed of protein family identifiers from a controlled vocabulary (Pfam, CAZy). We then inferred a probabilistic topic model (**2**). The topic variables of the model can be interpreted as potential functional modules, i.e. sets of functionally coupled protein families [43]. We obtained 400 modules with diverse biochemical functions. Next, we defined ‘genome-specific’ weights of the modules and used these weights in conjunction with the phenotype labels to rank the modules according to their estimated relevance for the phenotype of lignocellulose degradation (**3**). As weights, we used the fraction of protein families in a module that were present in a certain genome or metagenome bin (‘completeness scores’). We identified stable, high-ranking modules from independent repetitions of the analysis and constructed consensus modules, which we named plant biomass degradation modules (PDMs) (**4**). These PDMs were found to cover different aspects of plant biomass degradation, such as cellulose, hemicellulose and pectin degradation. Moreover, the weights of the PDMs could be used to predict the biomass degradation abilities of organisms, and we were able to identify specific gene clusters in the input set of (meta-)genomes which reflected the protein family content of individual modules (**5**). The clusters thus provided evidence for the functional coherence of the modules by gene neighborhood.

## Identification of stable plant biomass degradation modules (PDMs)

We used LDA based on Gibbs sampling for the inference of the modules, a Markov Chain Monte Carlo (MCMC) method which efficiently estimates parameters for complex models such as LDA. In agreement with the recommended procedures for MCMC sampling [47], we repeated the analysis multiple times (18 LDA runs) to ensure the stability of the results. We thus repeated the two central steps of our method, i.e. the inference of modules and their subsequent ranking by phenotype association (Figure 1, Steps 2 and 3), 18 times to identify stable, high-ranking modules. We summarized the information from stable, high-ranking modules found in different runs by constructing ‘consensus modules’ which contained all the protein families that were found in similar modules in at least nine LDA runs (Figure 1, Step 4; Methods).

We identified five consensus modules (M1–M5), which we refer to as plant biomass degradation modules (PDMs) (Table 1, Additional File **3**). We mapped the CAZy families of these PDMs to essential activities in the degradation of plant cell wall material based on their EC numbers (Table 2). All PDMs included protein families with cellulase-or hemicellulase activities, which supports the modules’ relevance for plant biomass degradation. M1–M5 were functionally distinct, with only a moderate overlap (12.6%) of their protein family content, including the broadly defined families GH5 and GH43 [48]. Isofunctional Pfam and CAZy terms, such as PF00150 and GH5, were grouped together into the same PDMs in most cases. The modules also included 20 Pfam families without a direct link to plant biomass degradation, such as ‘domains of unknown function’ (DUFs), ‘ricin-type β-trefoil lectin-like domains’ and ‘GDSL-like lipase/acylhydrolase’ (Table 3, Section 1 of Supplementary Note in Additional File **2**). Some of these domains could encode novel functions that are important for plant biomass degradation.

**Table 1:**
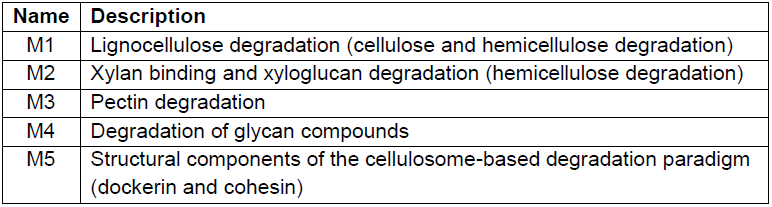
Functional characterization of the consensus plant biomass degradation modules M1–M5. We characterized each module based on the set of protein families contained in it. Additional File **3** shows each consensus module as a list of Pfam/CAZy terms.

| Name | Description |
| --- | --- |
| M1 | Lignocellulose degradation (cellulose and hemicellulose degradation) |
| M2 | Xylan binding and xyloglucan degradation (hemicellulose degradation) |
| M3 | Pectin degradation |
| M4 | Degradation of glycan compounds |
| M5 | Structural components of the cellulosome-based degradation paradigm (dockerin and cohesin) |

**Table 2:**
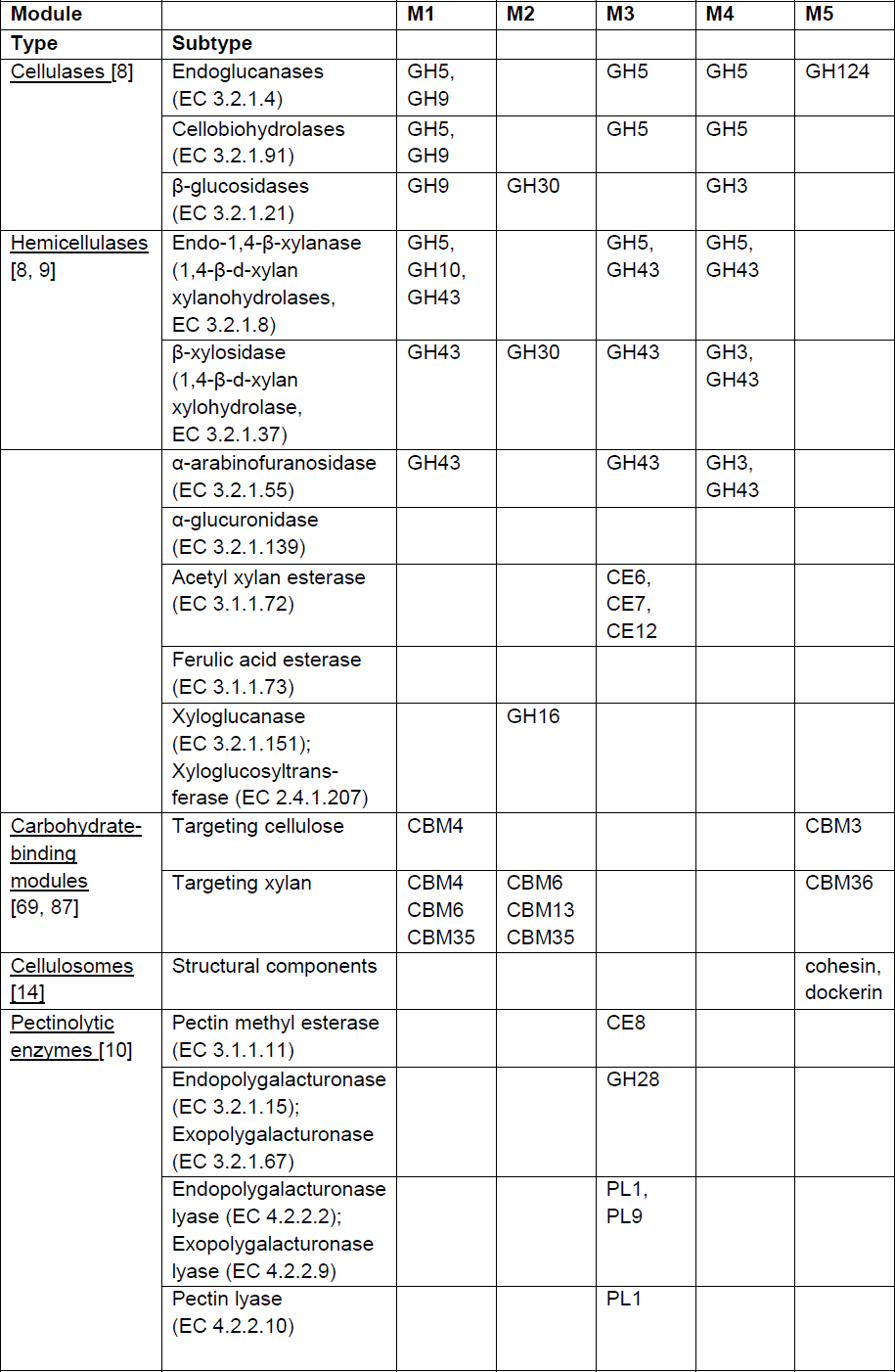
CAZy families in the PDMs M1–M5 with key functions for plant cell wall degradation.

**Table 3:**
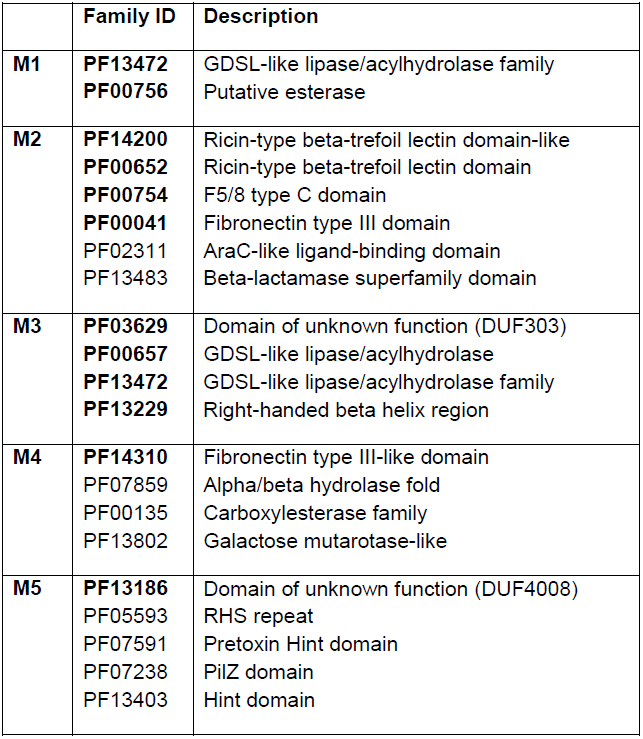
Protein families of the PDMs M1–M5 with potential functions in plant biomass degradation. Protein families that appeared in the gene clusters identified by mapping the PDMs to the phenotype(+) genomes are marked in bold.

## Gene clusters with PDM protein families

To confirm a functional context for the protein families assigned to the same PDM, we searched for gene clusters annotated with multiple families of a PDM in the 38 genomes of known degraders, as the proximity of genes within a genome indicates a shared functional context [49, 50] (Figure 1, Step 5). For each PDM, we identified gene clusters of four or more neighboring genes, with intergenic distances of ≤ 2 kb between consecutive genes. Overall, 81 gene clusters were found for the five PDMs, which represented 51 distinct, non-overlapping clusters. On average, 70.7% of the family content of each PDM could be mapped to gene clusters in known degraders. Some of the gene clusters discovered have been described as being active in lignocellulose degradation (see following sections), whereas the novel ones are candidates for further experimental investigation. Notably, eleven of the 20 protein families that have no direct link to lignocellulose degradation (Table 3) appeared in at least one gene cluster identified in known degrading species, which supports their potential role in the degradation process.

## Assessment of the PDMs’ potentials to predict unknown lignocellulose degraders

The completeness of a PDM in a genome was predictive for the ability of an organism to degrade lignocellulosic plant biomass. We determined the predictive value for each PDM in standard evaluation protocols with leave-one-out (LOO) and tenfold cross-validation experiments (Methods). In these experiments, genomes from the learning set of 120 known lignocellulose degraders and non-degraders were successively left out of the process of determining the completeness threshold. Subsequently, PDMs were predicted to be present in the omitted genomes if their completeness score for the genome was equal to or above the inferred threshold. This procedure was used to assess the generalization error of a PDM-based classifier to avoid overly optimistic performance estimates [51, 52]. We observed high F-scores for the PDMs in the LOO setting (82.1–96.2%) and lower bounds for the cross-validation estimates of prediction accuracy between 76.57% and 91.69% (Table 4).

**Table 4:**
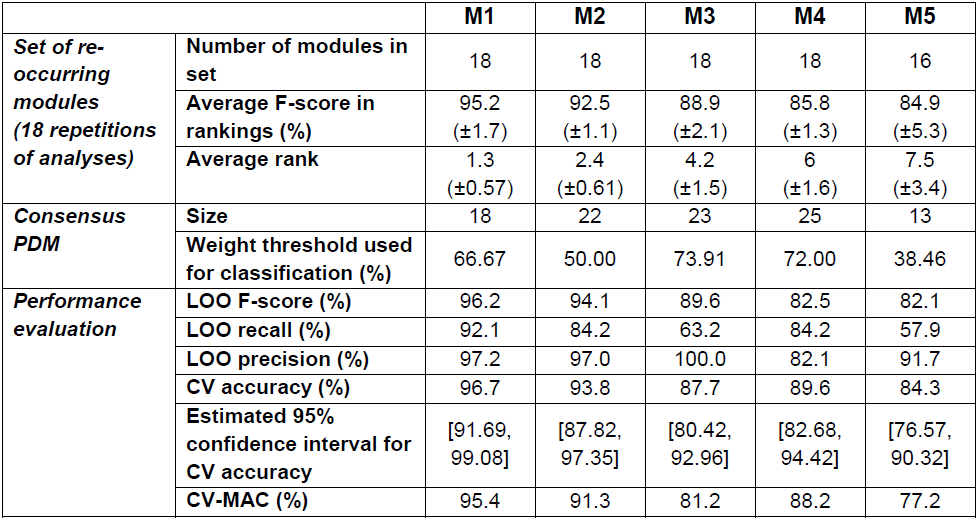
Association with lignocellulose degradation based on different performance measures for the consensus PDMs M1–M5. Each consensus PDM represents a set of re-occurring modules from 18 independent repetitions of our analysis (Figure 1) and contains all families that occurred in at least nine of these modules. The modules used to build the PDMs were identified by finding modules having minimal pairwise distances from each other (Methods). We report the average rank and average F-score of these module sets. ‘Size’ gives the number of Pfam and/or CAZy families that are contained in a PDM. We computed recall, precision and the F-measure scores for the individual PDMs in leave-one out (LOO) validation. In addition, accuracies and estimated confidence intervals for tenfold cross-validation (CV) are given to assess the generalization error more accurately. Following [28], we computed the cross-validation macro-accuracy (CV-MAC) as the average of the true positive and true negative rates.

|  |  | **M1** | **M2** | **M3** | **M4** | **M5** |
| --- | --- | --- | --- | --- | --- | --- |
| **Set of re-occurring modules (18 repetitions of analyses)** | **Number of modules in set** | 18 | 18 | 18 | 18 | 16 |
| | **Average F-score in rankings (%)** | 95.2<br>($\pm 1.7$) | 92.5<br>($\pm 1.1$) | 88.9<br>($\pm 2.1$) | 85.8<br>($\pm 1.3$) | 84.9<br>($\pm 5.3$) |
| | **Average rank** | 1.3<br>($\pm 0.57$) | 2.4<br>($\pm 0.61$) | 4.2<br>($\pm 1.5$) | 6<br>($\pm 1.6$) | 7.5<br>($\pm 3.4$) |
| **Consensus PDM** | **Size** | 18 | 22 | 23 | 25 | 13 |
|  | **Weight threshold used for classification (%)** | 66.67 | 50.00 | 73.91 | 72.00 | 38.46 |
| **Performance evaluation** | **LOO F-score (%)** | 96.2 | 94.1 | 89.6 | 82.5 | 82.1 |
|  | **LOO recall (%)** | 92.1 | 84.2 | 63.2 | 84.2 | 57.9 |
|  | **LOO precision (%)** | 97.2 | 97.0 | 100.0 | 82.1 | 91.7 |
|  | **CV accuracy (%)** | 96.7 | 93.8 | 87.7 | 89.6 | 84.3 |
|  | **Estimated 95% confidence interval for CV accuracy** | [91.69, 99.08] | [87.82, 97.35] | [80.42, 92.96] | [82.68, 94.42] | [76.57, 90.32] |
|  | **CV-MAC (%)** | 95.4 | 91.3 | 81.2 | 88.2 | 77.2 |

The top-ranking PDMs, M1 and M2, predicted the ability to degrade lignocellulose with cross-validation accuracies of more than 93%. Four genomes were misclassified by both M1 and M2 (figure in Additional File **4**, Tables in Additional File **7**): *Bryantella formatexigens* (false negative (FN)), *Xylanimonas cellulosilytica* (FN), *Thermonospora curvata* 43183 (FN) and *Actinosynnema mirum* (false positive (FP)). Interestingly, *A. mirum* and *T. curvata* might have been mischaracterized before [53], supporting the predictions by the two PDMs (Section 2 of Supplementary Note). All PDMs showed a precision of more than 82% for lignocellulose degraders, with few occurrences predicted for non-degraders. M3 and M5 were found only in a subset of the known degraders (Table 4: the lowest recall was 57.9%), suggesting that they might represent specific aspects of degradation strategies. However, looking at the presence/absence profiles of the PDMs across the degrading species, none of the PDMs showed an exclusive association with a known degradation paradigm (Figure 2).

**Figure 2:**
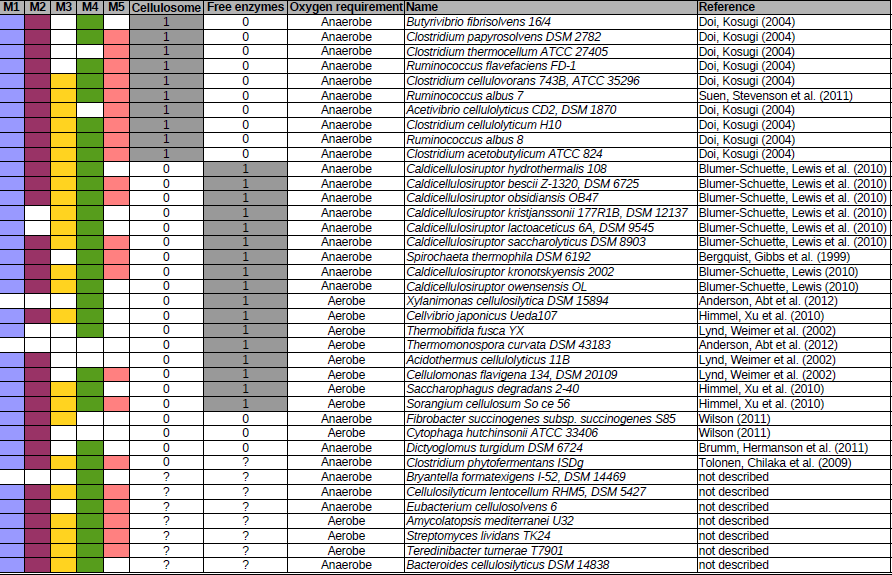
Occurrences of PDMs in organisms using different degradation paradigms. The predicted occurrences of the PDMs M1–M5 in the genomes of 38 known lignocellulose degraders are indicated by different colors. Each PDM was predicted to be present or absent from a genome, depending on its ‘genome-specific’ weight, i.e. the degree of completeness for its protein families. Two major cellulose degradation paradigms, the free enzyme and cellulosome-based strategies, were assigned to the organisms according to the literature. Assignments can be ambiguous; for example, *C. thermocellum* seems to be able to use mixed strategies [48]. No PDM was exclusively associated with these two paradigms, including M5, which, in addition to the cohesin and dockerin domains of cellulosomes, also included non-cellulosomal protein families (Table 3).

## Protein families of the PDMs

The highest-scoring PDM M1 (F-measure: 96.2%) incorporated various key families for the degradation of cellulose and hemicelluloses (Table 2), namely GH5, GH9, GH10, GH26, GH43 and CBM6 [48]. The GH5 and GH9 families together cover three classes of important cellulases [8]: Endoglucanases, cellobiohydrolases and β-glucosidases. Both are large families of cellulases which have been studied in many lignocellulolytic organisms (Section 3 of Supplementary Note). In addition to their cellulase activities, some members of these families are also hemicellulases with characterized activity on β-glucans, xyloglucans and heteroxylans [11]. The GH10 and GH43 families include xylanases and arabinases. M1 was present almost exclusively in lignocellulose-degrading bacteria (97.2% precision) and in almost all of them (92.1% recall). Similarly, also the individual modules used for creating the M1 consensus PDM showed strong associations with plant biomass degradation: M1 was always among the top three modules and was the top-ranked module in 14 of 18 LDA runs.

M2 (F-measure: 94.1%) contained families that bind and degrade xylan, xyloglucan and β-glucan (Table 2), such as GH30 (β-xylosidases), GH16 (β-glucanases, xyloglucanases) [9], CBM61 (which is often found with GH16) and the fucose-binding module CBM47. In addition, M2 included the xylan-binding domains CBM6, CBM35 and PF02018, which were also present in hemicellulolytic gene clusters with M2 families of *Clostridium cellulolyticum* and *Fibrobacter succinogenes* (Figure 3B, figure in Additional File **5**). In *Streptomyces lividans,* several small gene clusters of two or three genes with M2 member families might be linked to a xylan-binding mechanism involving CBM13 (also known as the ‘ricin superfamily’ or ‘R-type lectins’) [54]. CBM13 and two ‘ricin-type-β-trefoil lectin’ domains (PF14200 and PF00652 in Table 3) belonged to M2 and occurred in the clusters. Interestingly, the two different functional aspects of M2, i.e. xyloglucan degradation and xylan binding, were reflected by a split of the M2 module into two modules in some LDA runs.

**Figure 3:**
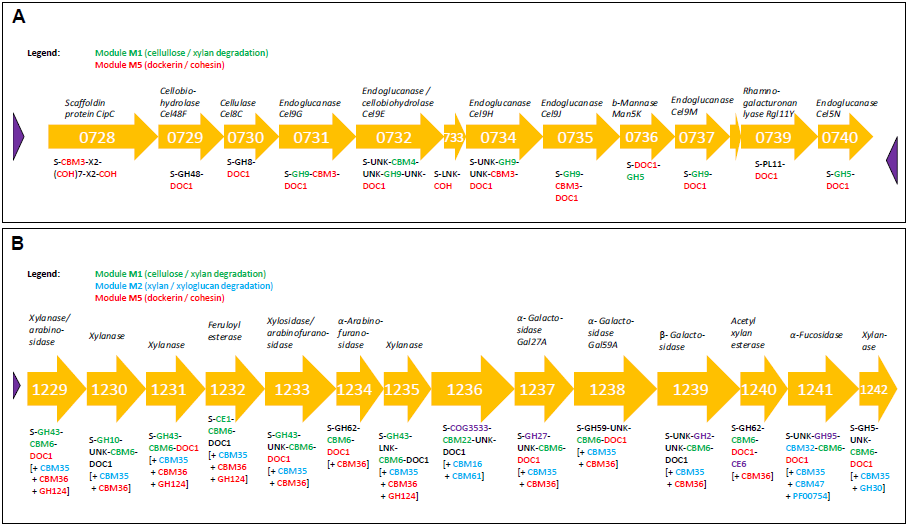
PDMs mapping to the *cel-cip* and *xyl-doc* gene clusters in *Clostridium cellulolyticus* H10. Blouzard *et al.* described two clusters of genes that are involved in cellulose and hemicellulose degradation [60]. Their domain architecture was adopted. Abbreviations used for the carbohydrate-binding module (CBM) and glycoside hydrolase (GH) architecture are: S: signal sequence; DOC1: dockerin type-I module; COH: cohesin type-I module; LNK: linker sequence; UNK: unknown function. We marked additional predicted domains as part of our in-house annotation sets using [+ family X]. Some dockerin annotations were filtered out by our bit score criterion. **Panel A:** Genes from the *cel-cip* operon (Ccel_0728 to Ccel_0740) are essential for the cellulose degradation ability of the organism *C. cellulolyticus* H10, which uses the cellulosome strategy. The cluster includes multiple protein families of the PDMs M1 and M5. Although the consensus modules of M1 and M5 did not directly include the two endoglucanase families GH8 and GH48, associations between M1 and GH8, and between M5 and GH48 existed (topic-word probabilities ≥0.005). **Panel B:** Genes from the 32-kb *xyl-doc* gene cluster (Ccel_1229 to Ccel_1242) encode functionalities for hemicellulose degradation. The cluster includes multiple protein families of the PDMs M1, M2 and M5, which together cover most of the cluster. Some additional protein families originate from M3 and M4 (purple). The following correspondences were used: CE1 ∼ PF00756 (esterase); CBM22 ∼ PF02018, and COG3533 (an uncharacterized protein in bacteria) ∼ PF07944 (a putative glycosyl hydrolase of unknown function, DUF1680). The *xyl-doc* cluster contains a xylosidase/arabinofuranosidase gene (Ccel_1233), which is characterized as a ‘putative β-xylosidase’ in IMG. The gene corresponds to β*-*xylosidase genes in *Caldicellulosiruptor saccharolyticus* (Csac_2411), *Bacteroides cellulosilyticus* (BACCELL_02584 and BACCELL_00858), and *Fibrobacter succinogenes* (FSU_2269/ Fisuc_1769). Clusters containing M1 protein families were also detected around these genes.

M3 (F-measure: 89.6%) included cellulose-degrading, hemicellulose-degrading and multiple pectinolytic enzymes (Table 2), such as pectin methyl esterase (CE8), pectin lyases PL1, PL9 and PF12708 (PL3) and endopolygalacturonase (GH28) (Table 2). M3 also included GH106 (α-L-rhamnosidase), which catalyzes the release of L-rhamnose from pectin (rhamnogalacturonan) molecules, and GH105, an unsaturated rhamnogalacturonyl hydrolase. Moreover, three acetyl xylan esterases (CE6, CE7 and CE12) were assigned to M3, as well as the uncharacterized domain PF03629 (DUF303), which may be an acetyl xylan esterase-related enzyme (InterPro accession: IPR005181). As CE12 has both acetyl xylan esterase (EC 3.1.1.72) and pectin acetylesterase (EC 3.1.1.-) activities assigned in CAZy, the other families are possibly also relevant for pectin degradation. Overall, the presence of multiple families involved in cellulose, hemicellulose and pectin degradation confirmed M3’s relevance for plant biomass degradation.

Module M4 (F-measure: 82.5%) contained the GH5, GH43, GH2 and GH3 families, as well as some associated Pfam domains, such as a GH2 sugar-binding domain (PF02837), and *C*- and *N*-terminal domains of GH3 (PF01915, PF00933). M4 also included GH35 and GH42, which are both β-galactosidases, and three members of a superfamily of α-galactosidases. D-galactose is an abundant component of the side chains of pectin, heteromannan and xyloglucan [7]. Activities in the degradation of pectins have been described for several β-galactosidases from plants [55]. Furthermore, M4 seemed to be linked to xyloglucan degradation in *B. cellulosilyticus* and C. *japonicus* (Section 4 of Supplementary Note). In conclusion, M4 comprised functionally diverse glycan degradation families, in line with the heterogeneous nature of hemicellulose polysaccharides [7] and their widely varying constituent sugars.

M5 (F-measure: 82.1%) included structural components of the cellulosome complex (cohesin and dockerin domains), the endoglucanase family GH124, and CBMs targeting cellulose (CBM3) and hemicellulose (CBM36). CBM3 is frequently found as a domain of cellulosome scaffoldin proteins [14]. The S-layer homology domain (PF00395), which anchors cellulosomes to the bacterial cell surface [14], was not associated with M5. It was consistently grouped into modules without significant scores in our rankings, indicating that the S-layer homology domain could perform other functions in non-degraders. M5 included five more Pfam domains of unknown relevance which are interesting candidates for novel functional activities (Table 3). PF13186, a domain of unknown function in our dataset, was annotated for the gene Cthe_3076 in *C. thermocellum,* which lies directly upstream of a gene cluster (Cthe_3077–3080) that is responsible for the structural organization of the cellulosome [56]. However, PF13186 was also annotated for non-degrading genomes (Figure 4) and was referred to as an ‘iron-sulfur cluster-binding domain’ in a recently updated version of the Pfam database.

**Figure 4:**
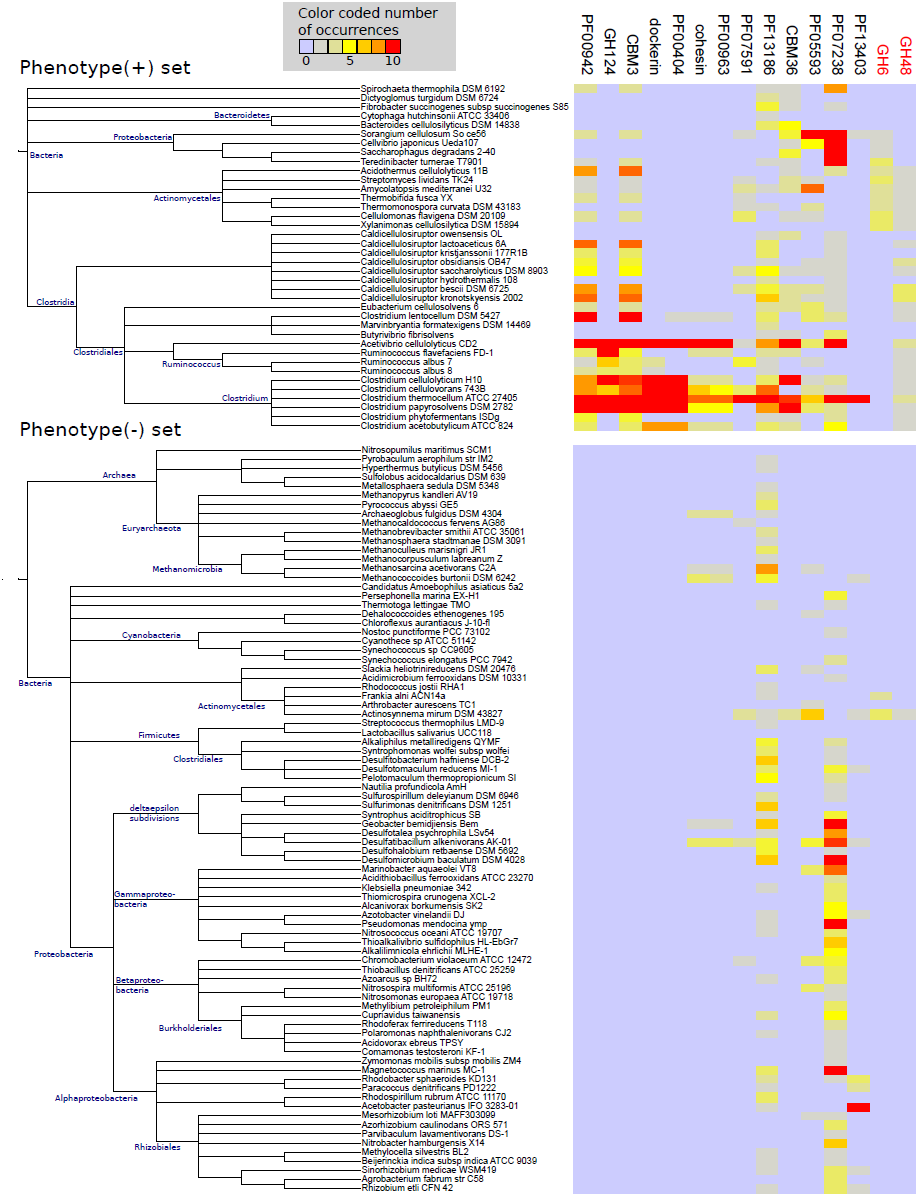
Co-occurrences of the M5 protein families and GH6, GH48 across known degraders and non-degraders. Combined co-occurrence profiles for the M5 protein families and two additional cellulases, GH6 and GH48, across the known sets of the phenotype(+) and phenotype(−) genomes, respectively. GH6 and GH48 were not assigned into the PDM M5; in case of GH48 only because of our strict cutoff criteria. However, GH48 was weakly associated with M5 and belonged to the top 50 families of the majority of M5 modules that were used to construct the consensus module. The colors of the heat map cells represent the number of instances of each family in the respective genomes of the organisms (see legends and note that the counted number of instances was limited to a maximum of ten per genome, as described in the Methods). The phylogenetic relationships of the genomes are indicated by dendrograms alongside the rows of the heat maps.

The protein families of the consensus PDMs are given in Additional File **3**. The file also lists all the less strongly associated PDM families that were found in fewer than nine similar modules of the 18 LDA runs and were thus not included in the consensus PDMs.

## Absence of the cellulase families GH6 and GH48

Interestingly, none of the PDMs contained the cellulase families GH6 or GH48. Both play an important role in cellulose degradation in some bacteria, but are not universally found in known lignocellulose degraders. They were not identified in *Fibrobacter succinogenes*, *Cytophaga hutchinsonii* or several gut and rumen metagenomes with lignocellulose-degrading capabilities [17, 20, 24, 57]. In line with these findings, we found GH6 and GH48 to be annotated in less than 5% of the samples of our input collection, and only a single GH6 annotation (no GH48) in the metagenome bins. This rarity in our dataset caused weak co-occurrence signals and is likely the cause why both families were not assigned to the stable, high ranking modules (Section 5 of Supplementary Note). Despite of this, GH48 was among the top 50 protein families of ten functional modules used to derive the M5 consensus module. This association with M5 is in line with the fact that many bacterial cellulosomes include proteins from the GH48 family [58]. However, the probabilities for GH48 were less than the threshold value *C* = 0.01 that we required for inclusion into modules. This is also evident from a weaker co-occurrence of GH48 with the M5 protein families in lignocellulose degraders (Figure 4). Another family with rare occurrences was GH44 (endoglucanases and xyloglucanases [59]), which appeared in less than 2% of our data samples and was not grouped into any module. This family does not seem to be essential for all lignocellulose degraders, as its catalytic activities are also covered by the CAZy families GH5, GH9 and GH16 (Table 2) [11]. Overall, the observed rarity of GH6, GH44 and GH48 might indicate their non-universal nature across lignocellulose-degrading species. However, it might be possible that more remote homologs exist that were not identified with the current Pfam and CAZy models.

## PDMs mapping to known gene clusters of essential lignocellulose degradation genes

The gene clusters in known degrader genomes that were identified based on the protein families of the individual PDMs included well-characterized clusters of lignocellulolytic genes. For example, the modules M1 and M5 mapped to the *cip-cel* operon and the *xyl-doc* gene cluster in *Clostridium cellulolyticum* H10 (Figure 3). *Cip-cel* encodes genes that are essential for cellulose degradation; *xyl-doc* encodes hemicellulose degradation genes [60]. The genes from both clusters have a multi-domain architecture with catalytic and carbohydrate-binding domains [60]. Within M1, GH5, GH9 and CBM4 occurred in *cip-cel,* while CBM6, CBM35, GH10, GH43, PF00756 and PF02018 have been annotated for *xyl-doc*. Genes from both clusters also include the cohesin and dockerin domains, which reflects the cellulosome-based degradation paradigm used by *C. cellulolyticum* H10.

Interestingly, LDA assigned the cohesin and dockerin domains to the M5 module, despite of their co-occurrence with the M1 families in *cip-cel* and *xyl-doc*. This is probably due to the existence of M1 families in the genomes of organisms that do not have cellulosomes, such as *Thermobifida fusca,* which is a model organism for the ‘free enzyme’ paradigm (Section 6 of Supplementary Note). M1 also mapped to a hemicellulolytic gene cluster in *Fibrobacter succinogenes,* an organism without cellulosomes that uses an unknown degradation strategy [61, 62] (figure in Additional File **5**). Despite the evidence for a link between M5 and the cellulosome strategy, none of the PDMs proved to be exclusive for a particular degradation paradigm (Figure 2). As described above, the M5 module also contained five Pfam families whose functional descriptions have no known link to lignocellulose degradation (Table 3). These shared co-occurrence patterns with the cohesin and dockerin domains, but in contrast to these they also occurred in organisms using free cellulolytic enzymes, such as some *Caldicellulosiruptor* species (Figure 4). Thus, M5 also covered non-cellulosome related functionalities (Section 7 of Supplementary Note).

## Predicting the ability for plant biomass degradation

We predicted the presence of PDMs for the 3096 remaining genomes and taxonomic metagenome bins if their completeness was equal to or above the threshold determined for each PDM (Methods). Overall, the presence of one or more PDMs was predicted for 8.4% (28/332) of the taxonomic bins and 24.7% (683/2764) of the genomes (tables in Additional File **8**). Most genomes and bins to which M1 was assigned also had M2 assigned to them (82% of 132 M1 assignments occur jointly with M2 assignments). This agreed with the cellulose-and hemicellulose-degrading (M1) and hemicellulose-targeting (M2) enzymatic activities we determined for these modules, which are both essential for lignocellulose degradation [45]. The majority of all predictions, i.e. 52.5%, were exclusive to M4 (Venn diagram in Additional File **9**). As M4 included functionally diverse glycan degradation families and had the lowest precision (82.1%) of all modules for lignocellulose degraders, these assignments likely reflect a general ability of the respective organisms to degrade carbohydrate substrates of plant origin.

In a previous study [45], Medie *et al.* analyzed the distributions of CAZy families representing cellulases, hemicellulases and pectinases across ∼1500 complete bacterial genomes. They have classified almost 20% of these organisms as saprophytic bacteria, based on the presence of at least one cellulase and three or more hemicellulases or pectinases. Saprophytes feed on dead organic matter of plant origin and thus are likely to include lignocellulose-degrading species. Based on the same CAZy families and criteria as described in [45], we determined potential saprophytes in our dataset (Methods). In total, about one quarter (27.2%) of all 3216 genomes and metagenome bins fulfilled these. The genomes and metagenome bins with predicted PDM occurrences were enriched with potential saprophytes (75% of all predictions). This enrichment was particularly large for M1 (99%), M2 (91%) and M3 (100%). These results further support the notion that the ability to degrade plant biomass is a frequent trait of Bacteria and Archaea species.

The metagenome bins that were assigned PDMs came from cow rumen, reindeer rumen, manatee gut, Tammar wallaby gut and termite hindgut samples, and samples of a methylotrophic and a terephthalate-degrading community. Most of these communities, except the methylotrophic/terephthalate-degrading ones, are known to include lignocellulose-degrading community members; however, their taxonomic affiliations are only partly known [19, 63, 64]. The coverage and quality of protein coding sequences was heterogeneous across the 332 bins, which resulted in low numbers of protein family annotations for some of the bins. Sixty-three bins were annotated with less than ten protein family annotations, while the remaining bins contained 276 different protein families per bin on average. This probably explains why PDMs were predicted to be present in only 28 bins covering five major taxonomic clades (Figure 5). PDMs occurring in metagenome bins of Bacteroides, Prevotella and Lachnospiraceae (Clostridiales) were in line with the taxonomic affiliations of cellulose degraders found in mammalian gut and rumen microbial communities [65]. Furthermore, the PDMs accurately identified Bacteroidales and Treponema bins that have been shown to be involved in lignocellulose degradation in recent metagenome studies of the cow rumen [66] and termite hindgut [57], thus indicating the benefit of our method to guide the discovery of uncultured microbial taxa with lignocellulolytic activities. Our results also implied two archaeal extremophile species that have plant biomass degradation capabilities (Section 8 of Supplementary Note).

**Figure 5:**
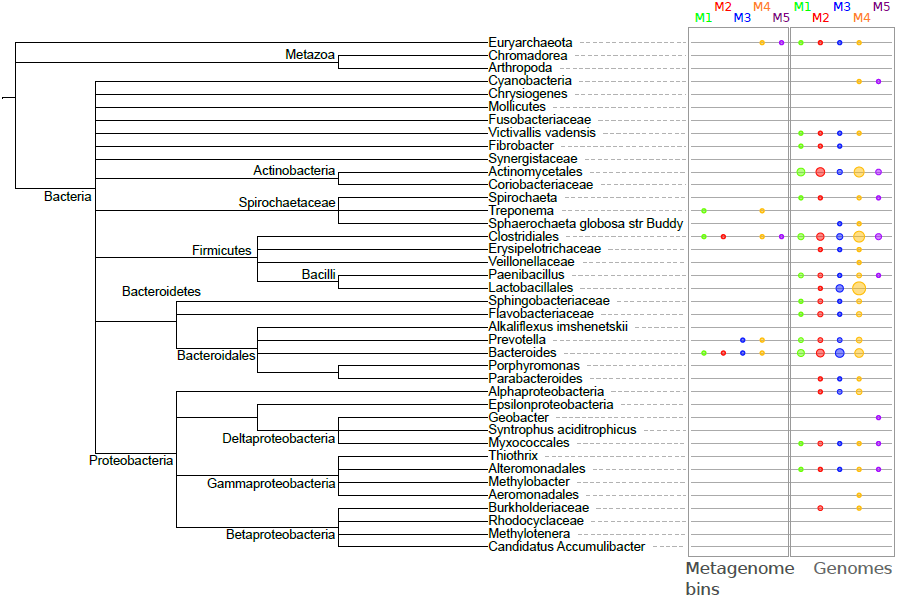
Comparison of PDM occurrences in metagenome bins and isolate genomes of corresponding taxa. The colored circles at the leaf nodes of the tree denote the predicted occurrences of the different PDMs in the respective taxa or their subclades. The tree was constructed from the taxonomic assignments of the metagenome bins in our input set (Methods). We then mapped the predicted occurrences of the PDMs in 28 different metagenome bins to the leaf nodes of the tree. If PDMs were identified in two or more bins, with one being a parental taxon to the other, the parent taxon is displayed. For example, predictions for *Prevotella ruminicola* were summarized with other predictions for the genus Prevotella. In addition, PDM occurrences in 403 isolate genomes of corresponding taxa were also mapped to the leaf nodes. The area of the colored circles for the isolate genomes was made proportional to the number of genomes for which the respective PDM was identified. The PDM predictions for the metagenome bins covered only five major taxonomic clades. The majority of PDMs in the metagenome bins were assigned to the orders Bacteroidales and Clostridiales. For the genomes, PDMs were identified from a broader range of taxa, including Actinobacteria, Firmicutes, Bacteroidetes and Proteobacteria, in agreement with the estimated taxonomic range of potential cellulose-degrading species reported in [46]. The differences in the identified PDMs and their taxonomic affiliations between genomes and metagenome bins may partly reflect the abundance of Bacteroidales and Clostridiales degraders in plant biomass-degrading communities, but some PDMs were likely also not identified in the metagenome bins due to the partial nature of the genomic information recovered.

## Identification of gene clusters and PULs in the predicted (meta)genomes

To identify new candidate clusters of genes encoding the ability to degrade lignocellulosic plant biomass, we searched for gene clusters encoding PDM protein families in the 711 genomes and taxonomic bins with PDMs, using the same criterion as before. We found 379 gene clusters of four or more genes for individual PDMs, which mapped to 342 distinct gene clusters in 168 genomes and six taxonomic bins. Genome fragmentation caused by incomplete assembly of bacterial draft genomes from IMG and taxonomic bins in our dataset may have decreased the number of detected clusters. Most of the gene clusters occurred in Bacteroidetes (54.3%); 22.4% and 12.7% occurred in Firmicutes and Actinobacteria, respectively. The first two phyla are predominant in gut and rumen environmental communities with lignocellulose-degrading abilities [65, 67]. Some of the newly identified gene clusters may cover polysaccharide utilization loci (PULs) targeting various kinds of polysaccharides. We found gene clusters in 39 isolate Bacteroides species, which are generally known to possess PULs [63]. As an example, the pectin-related PDM M3 identified gene clusters in *Bacteroides thetaiotaomicron* that represent parts of two regions that have been shown to be active in rhamnogalacturonan degradation in a PUL-targeted study [68]. Moreover, LDA inferred a stable functional module related to PULs which included a suite of outer membrane proteins as well as the two core proteins which are known to define PULs, namely SusD-and SusC(TonB)-like membrane proteins (Additional File **3**). This module was not part of the high-ranking modules, which can be explained by the broad substrate specificity of PULs for various polysaccharides, including starch in particular [15, 69]. While analyzing gene clusters of PDM protein families, we found hybrid gene clusters linking the PUL module to the glycoside hydrolases involved in lignocellulose degradation. For example, we identified gene clusters corresponding to previously characterized Sus-like PULs from *Bacteroides ovatus* targeting xyloglucan and xylan [68] (Section 9 of Supplementary Note).

## Predicting the ability for plant biomass degradation in a cow rumen microbial community

Hess *et al.* [19] reconstructed 15 draft genomes from the metagenome of a switchgrass-degrading microbial community from a cow rumen. Earlier, we cross-linked data from the cellulolytic enzyme screens of the original study with annotations of these draft genomes to identify plant biomass degraders [28]. Strikingly, the (hemi-)cellulolytic enzymes of the cow rumen bins with degradation abilities confirmed by activity screens (GH5, GH9, GH10 and GH26) were all part of M1 (Table 3 in [28]). We investigated whether PDM assignments allowed the identification of the plant biomass-degrading community members in the cow rumen metagenome (Table 5). The presence of M1 or M2 identified all degraders, in agreement with the enzyme screen results and our previous assignments with a family-centric SVM classifier [28]. M1 was also present in the draft genome ‘APb’, for which no lignocellulolytic enzymes were identified, but which is closely related to a known plant biomass-degrading species (*Butyrivibrio fibrisolvens*). The PDMs mapped to six gene clusters with four or more genes and several shorter clusters in the draft genomes. We investigated these and found an interesting cluster in the Bacteroidales-associated draft genome ‘AGa’ containing genes annotated with GH5, GH94 and two unannotated gene sequences (Section 10 of Supplementary Note).

**Table 5:** PDM assignments to draft genomes with the ability to degrade plant biomass within the cow rumen metagenome sample. Draft genomes with evidence for lignocellulolytic activity according to carbohydrolytic activity tests are indicated in green [19]. ‘APb’ was mapped using 16S RNA marker genes to the known lignocellulose-degrading organism *Butyrivibrio fibrisolvens* [19]. The draft genomes marked in bold were also predicted by a Support Vector Machine-based method for predicting lignocellulose degraders (counting only the unambiguous predictions of the SVM-classifier) [28].

| **Draft genome** | **Taxonomic affiliation** | **M1** | **M2** | **M3** | **M4** | **M5** |
| --- | --- | --- | --- | --- | --- | --- |
| **AJ** | Bacteroidales | √ | √ | √ | √ |  |
| **AGa** | Bacteroidales | √ |  |  |  |  |
| **AC2a** | Bacteroidales |  | √ |  |  |  |
| **Ala** | Clostridiales | √ |  |  | √ |  |
| **APb** | Clostridiales | √ |  |  | √ |  |
| **AH** | Bacteroidales | √ |  |  | √ |  |
| AFa | Spirochaetales |  |  |  | √ |  |
| AN | Clostridiales |  |  |  | √ |  |
| AWa | Clostridiales |  |  |  | √ |  |
| ATa | Clostridiales |  |  |  |  | √ |
| ADa | Myxococcales |  |  |  |  |  |
| AMa | Spirochaetales |  |  |  |  |  |
| AQ | Bacteroidales |  |  |  |  |  |
| AS1a | Clostridiales |  |  |  |  |  |
| BOa | Clostridiales |  |  |  |  |  |

## Conclusions

The degradation of lignocellulosic plant biomass is a complex biological process with mechanisms across different microbial species that are currently only partially understood. We describe functional modules of protein families linked to plant biomass degradation, which we identified based on co-occurrence patterns and partial phenotype information. Using LDA, a state-of-the-art Bayesian inference method, we inferred 400 potential modules from a set of 2884 genomes and 332 taxonomic bins from 18 metagenomes. Such modules represent sets of functionally coupled protein families and cover a broad range of biochemical processes, as shown in [43]. We then determined the presence of modules in genomes of known lignocellulose-degrading species and non-degraders to calculate a ranking of the modules that reflected the strength of their association with the plant biomass degradation phenotype. We analyzed the stability of the top ranking modules across several executions of the LDA method and determined five consensus functional modules (PDMs) involved in plant biomass degradation.

Our approach allowed us to include genomes and taxonomic metagenome bins lacking phenotype information in the inference step, which largely expanded the dataset in comparison to our previous study, where we linked individual protein families to plant biomass degradation [28]. We included the learning set of 19 known bacterial lignocellulose degraders from that study and expanded it to 38 species overall. Despite this, the fraction of confirmed degrader species is still small, compared to the estimated abundance of potential plant biomass-degrading species reported in two other studies [45, 46]. Based on these estimates, 20–25% of our genome collection could possess plant biomass degradation capabilities. Our approach allowed us to also include data from these potential degraders into the inference of functional modules, together with metagenome data from plant biomass-degrading communities. To our knowledge, this was the first study that globally analyzed the available genome sequence and phenotype data to determine the functional modules of the protein families that are linked to plant biomass degradation.

Overall, the PDMs included many known protein families for the degradation of cellulose, xylan, xyloglucan and pectins, which are the main components of plant cell walls, with families targeting the same macromolecules being grouped together. In addition to those known families, our results indicate that 20% of the PDMs’ member families seem to be linked to plant biomass degradation, although their individual functions are less clear or unknown (Table 3). Even more potentially interesting families were found in the high-ranking modules but were not included in the consensus modules because they occurred in less than half of the modules used to construct the consensus. Some of these families might be interesting for further investigation. The functional coherence of PDM member families was also supported by their localization in gene clusters in lignocellulolytic microbes. This included several known clusters of lignocellulolytic enzymes, such as *cip-cel* and *xyl-doc* from *Clostridium cellulolyticum* H10. Based on the modules, we overall identified more than 400 gene clusters in our dataset, some of which covered known PULs targeting different kinds of polysaccharides.

Moreover, we investigated whether certain modules were specific to different degradation paradigms, as the module M5, for example, contained cellulosome-related families, such as cohesin, dockerin and CBM3. None of the modules was exclusive for a specific degradation strategy, and the modules instead spanned different paradigms. We believe that the granularity of the modules could be further improved in the future if more and better curated phenotype information becomes available, which would allow us to enrich the set of genomes with species with different confirmed paradigms. For instance, the identification of genes from Sus-like cellulose-interacting protein complexes, as reported by Pope and Mackenzie [63], and Naas *et al*. [66], would likely require more accurate profile hidden Markov models for *susD*-like genes. For these, one would need to know the sequences of relevant genes in more organisms that use the Sus-like paradigm. Within our learning set, only *Bacteroides cellulolyticus* uses a Sus-like strategy on hemicellulosic polysaccharides [70].

The PDMs allowed us to predict the ability of lignocellulose degradation with cross-validation accuracies of up to 96.7%, which we used to predict the ability to degrade plant biomass for all genomes and taxonomic bins with unknown degradation status in our dataset. The predicted degraders were clearly enriched with organisms that were likely to have a saprophytic lifestyle. For 15 draft genomes of a microbial community from a cow rumen, we confirmed the predictions by cross-linking to enzymes with demonstrated lignocellulolytic activities. In addition, the PDMs identified cellulolytic metagenome bins for several cellulolytic metagenome communities.

The PDMs contained many of the protein families that we had previously identified with a family-centric approach in a smaller set of 19 known lignocellulose degraders and three metagenomes, including CBM3, CBM4, CBM_4_9, CBM6, GH5, GH10, GH26, GH43, GH55, GH88 and GH95 [28]. Apart from that, differences in the results existed. For example, in [28], only a few pectin-related families were identified, i.e. PL1, GH88 and GH106; however, here, we identified an entire module of pectin-degrading families (PDM M3), which included these three families together with PL3, PL9, GH28, GH105, CE12 and additional related ones. Differences were also found for individual families. For example, the PDMs were linked to GH9, GH48, cohesin and dockerin, as well as elements of xylan binding, such as the CBM13 and lectin domains, which were not identified with the family-centric approach. On the other hand, GH6 and GH44 were not associated with the PDMs. These families occurred in less than 5% of the input genomes and metagenome bins and their co-occurrence patterns with other families were more subtle in our large data collection than in the smaller dataset analyzed previously, which suggests their lower relevance on a global scale. Both families appear to be non-essential, as GH6 has been noted to be absent in several known degraders as described, and the catalytic activities of GH44 are also represented by the families GH5, GH9 and GH16 (Table 2). In addition to differences in dataset sizes and composition, methodological differences between the two approaches were likely to be responsible for the differences observed and the additional relevant families that were included in the PDMs. Neither approach identified any gene families related to lignin degradation. This may be because lignin-related protein domains, except for the broadly defined peroxidase family PF00141, were largely missing from the Pfam and CAZy/dbCAN databases. Furthermore, reports of lignin decomposition have been dominated by fungi [71], and thus the corresponding mechanisms might have been under-represented in our bacterial and archaeal dataset.

We showed evidence for functional links of the protein families in the PDMs with each other and the plant biomass degradation phenotype, which includes the co-occurrences of these families across genomes, co-occurrences with known relevant families, clustering within the genomes of known degraders and the predictive value of the PDMs for identifying plant biomass degraders. Given this extensive support, an experimental characterization of the PDMs’ protein families with unknown relevance for plant biomass degradation and their respective gene clusters is likely to reveal new biochemical functionalities for plant biomass degradation. With the method we have described, other phenotypes such as nitrogen fixation or antibiotic resistance could be studied from existing genome datasets in a similar fashion.

## Methods

### Latent Dirichlet Allocation

LDA is a text-mining method for extracting semantic concepts (i.e. topics) from a collection of text documents [42]. The topics reflect groups of semantically related words supported by co-occurrence signals across the document collection. LDA is a generative probabilistic model assuming a well-defined process as the source of the observed documents. With Bayesian inference and MCMC methods such as Gibbs sampling, the generative process can be inversed [72, 73]. This corresponds to increasing the model’s probability by fitting latent variables such that the outcome of the process matches the observed documents as closely as possible. Here, we are interested in inferring the latent variables, not the outcome of the process itself.

The input for LDA is a collection of *N* documents, where each document is a collection of words stemming from a controlled vocabulary *V*. The order of words in a document is not important, which is called the ‘bag of words’ assumption. LDA assumes the existence of *T* latent topics, and each topic is represented as a discrete multinomial distribution over *V.*

One variable of the model with central meaning is the vector 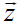, which contains a random variable *z* for each word of the text collection that models the word’s latent origin with respect to the *T* topics. According to the model, the probability of observing word *w* in document *d* of the collection is given by:

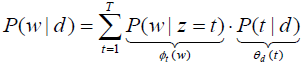

Here, 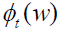 defines the multinomial distribution representing topic *t* and 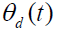 corresponds to a multinomial distribution describing the document-specific prior probabilities of the topics. The parameters 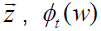 and 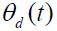 for all documents and topics are latent variables of the hidden generative process, which can be estimated quite efficiently with MCMC sampling methods.

## Genome and metagenome annotation

Protein sequences for bacterial and archaeal species were downloaded from IMG (version 3.4) and metagenomic protein sequences were obtained from ‘IMG with microbiome samples’ (IMG/M, version 3.3). In addition, samples of microbial communities from the Svalbard reindeer rumen [63], termite hindgut [57], manatee gut and the forestomach of the Tammar wallaby [64], as well as draft genomes reconstructed from a metagenome sample of a switchgrass-degrading microbial community in a cow rumen were included [19]. If no protein coding sequences were available, genes were predicted with MetaGeneMark [74]. Taxonomic bins from IMG/M or the original publications, or generated in-house, were used for all metagenome samples and were inferred with either *PhyloPythia* [75] or *PhyloPythiaS* [76] using sample-specific training sequences and taxonomic models constructed with taxa that represent the more abundant community populations. Overall, we worked with protein-coding sequences from 2884 prokaryotic genome sequences and 332 taxonomic bins derived from 18 metagenome samples. Protein sequences were annotated with protein families from Pfam (Pfam-A 26.0), CAZy [11] and dbCAN [77] using HMMER 3.0 [78]. Multiple matches of different domains per protein were allowed. Matches were required to have an e-value of at least 1e-02 and a bit score of 25 or more. Matches to the hidden Markov models of the CAZy families from dbCAN aligning for more than 100 bp with an e-value > 1e-04 were excluded. We then converted the protein domain annotations for the genomes and taxonomic metagenome bins into a suitable input collection for LDA (Section 1 of Supplementary Methods in Additional File **1**).

## Functional module inference with LDA

We used the protein family collection of the (meta-)genomes as input for the LDA inference procedure to predict potential functional modules, as demonstrated in [43]. Because of the larger input collection, we increased the number of topics from 200 to 400. Despite the increased number of documents compared to our previous work (3216 vs. 575), there was a slight decrease in the vocabulary size (8413 vs. 10,431), due to differences in coverage between the Pfam-A and eggNOG databases. As in [43], we used the parameter value *C* = *0.01* to convert topic probability distributions into discrete sets of protein domains, which represented our potential functional modules. Thus module *M_t_* for topic *t* was defined as 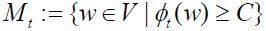 and contained the protein family identifiers that were most strongly related to topic *t.* The families assigned to *M_t_* share common co-occurrence patterns and were therefore likely to be functionally coupled due to the ‘guilt by association’ principle [79].

## Phenotype annotation

We assigned the lignocellulose-degrading phenotype to genomes by manually curating the annotations of ‘(ligno)cellulose degradation’ or ‘(plant) biomass degradation’ from IMG, the Genomes Online Database (GOLD) [80] and the German Collection of Microorganisms and Cell Cultures (DSMZ) (http://www.dsmz.de) based on information from the literature. Removal of ambiguous or inconsistent phenotype annotations resulted in 38 confirmed lignocellulose degraders (phenotype(+) genomes) which degraded some or all components of lignocellulose (table in Additional file **11**). The set of phenotype(+) genomes is a superset of the 19 lignocellulose-degrading microbes (except *Postia placenta*) from our previous work [28]. We adapted the set of 82 phenotype(−) genomes from the same study, which were also manually curated using information from the literature. There was less certainty in phenotype(−) annotations, as it may be that a particular phenotype has not been discussed in the literature; however, we used statistical methods to determine PDMs from these datasets that can tolerate a certain amount of error.

## Definition of module weights

The inference of a topic model with LDA from a collection of *N* input documents results in *T* potential functional modules. We extracted 400 modules from 3216 genomes and metagenome bins. We then applied an attribute ranking approach to sort the modules according to their relevance for lignocellulose degradation. As attributes to be used in the ranking procedure, we defined ‘module weights’. A weight, *weight_t_* (*d*), should reflect how likely the module *M_t_* is to be contained in the genome or metagenome bin encoded as document *d* of the input collection. Given *N* genomes or bins as input and *T* modules, we can summarize the weights in a weight matrix 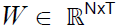 with entries 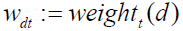.

Two different definitions of weights (probability weights and ‘completeness scores’) were tested (Section 2 of Supplementary Methods). We decided to use completeness scores, as they produced more relevant results, though the rankings obtained with both choices of weights largely agreed (Section 11 of Supplementary Note). The ‘completeness score’ of a module is the percentage of a module’s protein families that occurred in a specific genome or taxonomic bin. More precisely, we defined the weight of module *M_t_* in document *d* of the (meta-)genome collection based on completeness as:

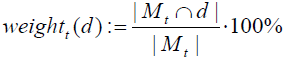

## Identification of phenotype-defining functional modules

To identify phenotype-associated modules, we used the weights of the modules in the input documents that corresponded to our manually curated phenotype(+) and phenotype(−) genomes. We refer to these genomes as the learning set. The selected weights were used to predict the phenotypes of these genomes and we scored each of the 400 modules according to its ability to distinguish between the two phenotype classes. More precisely, the classification of the learning set with respect to module *M_t_* was done by applying a threshold value *γ _t_* to the weights of the module, i.e. genomes were predicted to be phenotype(+) if the respective weights satisfied the threshold or phenotype(−) otherwise.

The ranking procedure optimized independent thresholds for all modules by finding the threshold that maximized a criterion function. We used the F-measure [67] with the parameter *β* = 0.5 for scoring (Section 3 of Supplementary Methods), which can be computed using the following confusion matrix:

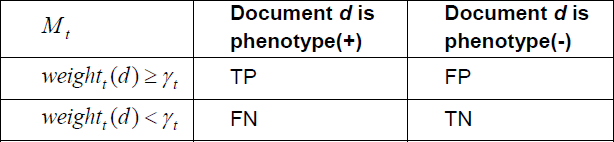

Finally, we obtained the ranking of the modules by sorting them in decreasing order based on their F-measure scores.

## Mapping of modules between Gibbs samples and runs

Finding the optimal assignments of protein families to functional modules, such that the observed data can be explained in the best possible way, is a combinatorially challenging task. We used Gibbs sampling to derive statistical estimates for the latent topic (functional module) distributions of the LDA model, from which we derived the potential functional modules as described. We then searched for similar modules across several LDA runs to identify stable modules, because with the MCMC inference technique used there is variance in the derived estimates across different runs. We used the Kullback–Leibler divergence [43] and the Jaccard distance [81] to calculate pairwise distances between topics (probability distributions) or modules (discrete protein family sets), respectively. As expected, we observed a good agreement between the results with both distance measures. Given the matrix of pairwise distances for the modules of two LDA runs, we used the Hungarian algorithm [82] to find an optimal global mapping between these. The Bron–Kerbosch algorithm [83] was used to find cliques of similar modules efficiently across multiple LDA runs (requiring pairwise KL-distances ≤ 5 or Jaccard distances ≤ 0.75, respectively).

## Consensus modules

In theory, Gibbs sampling efficiently estimates the posterior distribution of the model parameters and converges to a global optimum given a sufficient number of iterations [47]. However, in practice, we observe variance in the results of individual LDA runs and a common approach to derive a stable solution is to repeat the inference multiple times and to compare the results from a number of runs [72]. Therefore, we repeated the steps of our analysis several times with the same input data. In comparison to our previous study [43], we doubled the number of LDA runs to 18. In each run, we inferred 400 potential functional modules. As described in the previous section, we tracked the identities of the modules across all runs based on pairwise module distances and thus characterized the stability of the modules. Next, we applied the described attribute-ranking scheme based on the completeness scores to each of the 18 sets of 400 inferred modules and determined the top 15 modules for each run. Among these highly ranked modules from different runs, we searched for similar modules that occurred in at least 75% of the 18 runs. From these re-occurring modules we derived ‘consensus modules’ of protein families (Additional File **3**) as follows: Given a set of similar modules from different LDA runs, which were identified as representing a stable module across 75% or more of the 18 runs, the corresponding consensus module contains all protein families that occur in at least nine modules of this set.

## Leave-one-out analysis and tenfold cross-validation

For the consensus PDMs, we performed leave-one-out and tenfold cross-validation experiments to assess their predictive accuracy. In a loop, we successively left out each individual genome (or 10% of the genomes) of the learning set and optimized the weight threshold of a module on the remaining learning set with the F-measure. For the omitted genomes, the PDM was predicted to be present if the genome-specific module weight was equal to or above the inferred threshold. In both settings, we obtained exactly one prediction for each genome of the learning set, based on which we calculated performance measures such as precision and recall, the F-measure, the cross-validation accuracy and the cross-validation macro-accuracy. For the tenfold cross-validation experiments, we randomly split the data to create the different folds. The procedure was repeated ten times and the results were averaged. For a more accurate estimate of the test error, we also calculated 95% confidence intervals for the modules’ cross-validation accuracies. We used the Clopper–Pearson bound [52], which is an estimate based on the binomial distribution and the observed error rate on the omitted test samples. Note that the number of available test samples (120 in our case) is an important parameter of the binomial and determines the sizes of the intervals. With a larger set, one would obtain narrower bounds.

## Prediction of module occurrences in genomes and metagenome bins

We optimized the cutoff thresholds for module prediction by maximizing the F-measure using the weights of the consensus modules for all genomes with a known phenotype. We then considered the module weights in the genomes and metagenome bins of unknown phenotype to predict occurrences of the modules. We applied the following prediction rule to predict the presence of a module *M_t_* in the genome or metagenome bin *G_d_* that corresponds to document *d* in the input of LDA:

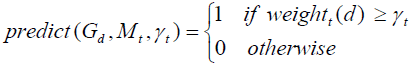

## Comparison of PDM occurrences in the taxonomic bins of metagenomes and isolate genomes of the corresponding clades

We constructed a tree based on the NCBI taxonomy tree with iTOL [84] for the taxa represented by the metagenome bins in the dataset. Metagenome bins with less than ten protein families were excluded from consideration. We used taxonomic assignments inferred by the binning methods *PhyloPythia* [75] and *PhyloPythiaS* [76], but not for the high-ranking bins, such as bacteria. To visualize the common occurrences of the PDMs at the leaf nodes of the tree, we collapsed some of the original leaf nodes to new leaf nodes of higher ranks. This was done if two or more of the PDMs were predicted to occur in taxa of the same clade, but with different ranks. In these cases, the PDMs involved were displayed for the highest common rank that was observed. The PDMs were predicted to occur in the bins of only five major taxa across the different metagenomes (Figure 5). In addition, we also mapped isolate genomes of the corresponding taxa with predicted occurrences of the PDMs to the leaf nodes of the tree.

## Identification of saprophytic genomes and taxonomic bins

As described by Medie *et al*. [45], we classified a genome or metagenome bin as belonging to a ‘cellulase-and hemicellulase containing saprophyte’ if the corresponding annotation set contained at least one cellulase and three or more hemicellulases or pectinases from the following families:

- *Cellulase families:* GH5, GH6, GH8, GH9, GH12, GH44, GH45, GH48, GH74 and GH124;
- *Hemicellulase and pectinase families:* GH10, GH11, GH26, GH28, GH30, GH43, GH53, GH67, GH78, PL1, PL2, PL9, PL10, PL11 and PL22.

## Implementation and parameter settings

We used the LDA implementation from the topic modeling toolbox (http://psiexp.ss.uci.edu/research/programs_data/toolbox.htm). The LDA model depends on two hyperparameters, *α* and *β*, which control the Dirichlet priors of the multinomial distributions. We used the default values of the topic modeling toolbox, i.e. *α* =50*/T* and *β* =0.01.

## List of abbreviations

CAZy: Carbohydrate-active enzyme
CBM: Carbohydrate-binding module
CE: Carbohydrate esterase
DUF: Domain of unknown function (in the Pfam database)
GH: Glycoside hydrolase
LDA: Latent Dirichlet Allocation
PDM: Plant biomass degradation module
PL: Polysaccharide lyase
PUL: Polysaccharide utilization locus
Sus: Starch utilization system

## Competing interests

The authors declare that they have no competing interests.

## Authors’ contributions

SGAK and ACM designed the study, interpreted the results and wrote the manuscript. SGAK conducted the experiments under the supervision of ACM. SGAK and AW computed the CAZy and Pfam protein annotations and curated the learning set. PBP was involved in the interpretation of the results and revised the final manuscript. All authors read and approved the manuscript.

## Acknowledgements

SGAK, AW and ACM were supported by the Max Planck society and Heinrich Heine University Düsseldorf. PBP gratefully acknowledges support from the Research Council of Norway (Project number 214042).

## References

1. Kumar R, Singh S, Singh OV: Bioconversion of lignocellulosic biomass: biochemical and molecular perspectives. J Ind Microbiol Biotechnol 2008, 35:377–391.

2. Kohse-Hoinghaus K, Osswald P, Cool TA, Kasper T, Hansen N, Qi F, Westbrook CK, Westmoreland PR: Biofuel combustion chemistry: From ethanol to biodiesel. Angew Chem Int Ed Engl 2010, 49:3572–3597.

3. Himmel ME, Ding SY, Johnson DK, Adney WS, Nimlos MR, Brady JW, Foust TD: Biomass recalcitrance: Engineering plants and enzymes for biofuels production. Science 2007, 315:804–807.

4. Gowen CM, Fong SS: Exploring biodiversity for cellulosic biofuel production. Chem Biodivers 2010, 7:1086–1097.

5. Xing MN, Zhang XZ, Huang H: Application of metagenomic techniques in mining enzymes from microbial communities for biofuel synthesis. Biotechnol Adv 2012, 30:920–929.

6. Minic Z, Jouanin L: Plant glycoside hydrolases involved in cell wall polysaccharide degradation. Plant Physiol Biochem 2006, 44:435–449.

7. Burton RA, Gidley MJ, Fincher GB: Heterogeneity in the chemistry, structure and function of plant cell walls. Nat Chem Biol 2010, 6:724–732.

8. Sweeney MD, Xu F: Biomass converting enzymes as industrial biocatalysts for fuels and chemicals: Recent developments. Catalysts 2012, 2:244–263.

9. Gilbert HJ, Stalbrand H, Brumer H: How the walls come crumbling down: Recent structural biochemistry of plant polysaccharide degradation. Curr Opin Plant Biol 2008, 11:338–348.

10. Jayani RS, Saxena S, Gupta R: Microbial pectinolytic enzymes: A review. Process Biochem 2005, 40:2931–2944.

11. Cantarel BL, Coutinho PM, Rancurel C, Bernard T, Lombard V, Henrissat B: The carbohydrate-active enzymes database (CAZy): An expert resource for glycogenomics. Nucleic Acids Res 2009, 37:D233–238.

12. Morais S, Barak Y, Lamed R, Wilson DB, Xu Q, Himmel ME, Bayer EA: Paradigmatic status of an endo-and exoglucanase and its effect on crystalline cellulose degradation. Biotechnol Biofuels 2012, 5:78.

13. Wilson DB: Microbial diversity of cellulose hydrolysis. Curr Opin Microbiol 2011, 14:259–263.

14. Fontes CM, Gilbert HJ: Cellulosomes: Highly efficient nanomachines designed to deconstruct plant cell wall complex carbohydrates. Annu Rev Biochem 2010, 79:655–681.

15. Martens EC, Koropatkin NM, Smith TJ, Gordon JI: Complex glycan catabolism by the human gut microbiota: The Bacteroidetes Sus-like paradigm. J Biol Chem 2009, 284:24673–24677.

16. Bolam DN, Koropatkin NM: Glycan recognition by the Bacteroidetes Sus-like systems. Curr Opin Struct Biol 2012, 22:563–569.

17. Wilson D: Evidence for a novel mechanism of microbial cellulose degradation. Cellulose 2009, 16:723–727.

18. Horn SJ, Vaaje-Kolstad G, Westereng B, Eijsink VG: Novel enzymes for the degradation of cellulose. Biotechnol Biofuels 2012, 5:45.

19. Hess M, Sczyrba A, Egan R, Kim TW, Chokhawala H, Schroth G, Luo S, Clark DS, Chen F, Zhang T, et al: Metagenomic discovery of biomass-degrading genes and genomes from cow rumen. Science 2011, 331:463–467.

20. Pope PB, Mackenzie AK, Gregor I, Smith W, Sundset MA, McHardy AC, Morrison M, Eijsink VG: Metagenomics of the Svalbard reindeer rumen microbiome reveals abundance of polysaccharide utilization loci. PLoS ONE 2012, 7:e38571.

21. Graham JE, Clark ME, Nadler DC, Huffer S, Chokhawala HA, Rowland SE, Blanch HW, Clark DS, Robb FT: Identification and characterization of a multidomain hyperthermophilic cellulase from an archaeal enrichment. Nat Commun 2011, 2:375.

22. Kim SJ, Lee CM, Han BR, Kim MY, Yeo YS, Yoon SH, Koo BS, Jun HK: Characterization of a gene encoding cellulase from uncultured soil bacteria. FEMS Microbiol Lett 2008, 282:44–51.

23. Wang F, Li F, Chen G, Liu W: Isolation and characterization of novel cellulase genes from uncultured microorganisms in different environmental niches. Microbiol Res 2009, 164:650–657.

24. Duan C-J, Feng J-X: Mining metagenomes for novel cellulase genes. Biotechnol Lett 2010, 32:1765–1775.

25. Rubin EM: Genomics of cellulosic biofuels. Nature 2008, 454:841–845.

26. Park BH, Karpinets TV, Syed MH, Leuze MR, Uberbacher EC: CAZymes Analysis Toolkit (CAT): Web service for searching and analyzing carbohydrate-active enzymes in a newly sequenced organism using CAZy database. Glycobiology 2010, 20:1574–1584.

27. Wang PI, Marcotte EM: It’s the machine that matters: Predicting gene function and phenotype from protein networks. J Proteomics 2010, 73:2277–2289.

28. Weimann A, Trukhina Y, Pope PB, Konietzny SG, McHardy AC: *De novo* prediction of the genomic components and capabilities for microbial plant biomass degradation from (meta-)genomes. Biotechnol Biofuels 2013, 6:24.

29. Slonim N, Elemento O, Tavazoie S: *Ab initio* genotype-phenotype association reveals intrinsic modularity in genetic networks. Mol Syst Biol 2006, 2:1–14.

30. Lingner T, Muhlhausen S, Gabaldon T, Notredame C, Meinicke P: Predicting phenotypic traits of prokaryotes from protein domain frequencies. BMC Bioinformatics 2010, 11:481.

31. Liu B, Pop M: MetaPath: Identifying differentially abundant metabolic pathways in metagenomic datasets. BMC Proc 2011, 5 Suppl 2:S9.

32. Kastenmuller G, Schenk ME, Gasteiger J, Mewes HW: Uncovering metabolic pathways relevant to phenotypic traits of microbial genomes. Genome Biol 2009, 10:R28.

33. Schmidt MC, Rocha AM, Padmanabhan K, Shpanskaya Y, Banfield J, Scott K, Mihelcic JR, Samatova NF: NIBBS-search for fast and accurate prediction of phenotype-biased metabolic systems. PLoS Comput Biol 2012, 8:e1002490.

34. Yosef N, Gramm J, Wang Q-F, Noble WS, Karp RM, Sharan R: Prediction of phenotype information from genotype data. Commun Inf Syst 2010, 10:99–114.

35. Padmanabhan K, Wilson K, Rocha AM, Wang K, Mihelcic JR, Samatova NF: In-silico identification of phenotype-biased functional modules. Proteome Sci 2012, 10 Suppl 1:S2.

36. Vey G, Moreno-Hagelsieb G: Metagenomic annotation networks: Construction and applications. PLoS ONE 2012, 7:e41283.

37. Jeffery C: Moonlighting proteins: Implications and complications for proteomics. Protein Sci 2004, 13:124–124.

38. Caetano-Anolles G, Yafremava LS, Gee H, Caetano-Anolles D, Kim HS, Mittenthal JE: The origin and evolution of modern metabolism. Int J Biochem Cell Biol 2009, 41:285–297.

39. De Filippo C, Ramazzotti M, Fontana P, Cavalieri D: Bioinformatic approaches for functional annotation and pathway inference in metagenomics data. Brief Bioinform 2012, 13:696–710.

40. Aravind L: Guilt by association: contextual information in genome analysis. Genome Res 2000, 10:1074–1077.

41. Goh CS, Bogan AA, Joachimiak M, Walther D, Cohen FE: Co-evolution of proteins with their interaction partners. J Mol Biol 2000, 299:283–293.

42. Blei DM, Ng AY, Jordan MI: Latent Dirichlet Allocation. J Mach Learn Res 2003, 3:993–1022.

43. Konietzny SG, Dietz L, McHardy AC: Inferring functional modules of protein families with probabilistic topic models. BMC Bioinformatics 2011, 12:141.

44. von Mering C, Jensen LJ, Snel B, Hooper SD, Krupp M, Foglierini M, Jouffre N, Huynen MA, Bork P: STRING: known and predicted protein-protein associations, integrated and transferred across organisms. Nucleic Acids Res 2005, 33:D433–437.

45. Medie FM, Davies GJ, Drancourt M, Henrissat B: Genome analyses highlight the different biological roles of cellulases. Nature Reviews Microbiology 2012, 10:227–234.

46. Berlemont R, Martiny AC: Phylogenetic distribution of potential cellulases in bacteria. Appl Environ Microbiol 2013, 79:1545–1554.

47. Gilks WR, Richardson S, Spiegelhalter DJ: Markov Chain Monte Carlo in Practice Boca Raton, Florida, USA: Chapman and Hall/CRC; 1999.

48. Himmel ME, Xu Q, Luo Y, Ding S-Y, Lamed R, Bayer EA: Microbial enzyme systems for biomass conversion: Emerging paradigms. Biofuels 2010, 1:323–341.

49. Overbeek R, Fonstein M, D’Souza M, Pusch GD, Maltsev N: The use of gene clusters to infer functional coupling. Proc Natl Acad Sci U S A 1999, 96:2896–2901.

50. Ballouz S, Francis AR, Lan R, Tanaka MM: Conditions for the evolution of gene clusters in bacterial genomes. PLoS Comput Biol 2010, 6:e1000672.

51. Duda RO, Hart PE, Stork DG: Pattern classification. John Wiley & Sons; 2012.

52. Anguita D, Ghelardoni L, Ghio A, Ridella S: Test error bounds for classifiers: A survey of old and new results. In Foundations of Computational Intelligence (FOCI), 2011 IEEE Symposium on; 11–15 April 2011. 80–87.

53. Anderson I, Abt B, Lykidis A, Klenk HP, Kyrpides N, Ivanova N: Genomics of aerobic cellulose utilization systems in actinobacteria. PLoS ONE 2012, 7:e39331.

54. Boraston AB, Tomme P, Amandoron EA, Kilburn DG: A novel mechanism of xylan binding by a lectin-like module from *Streptomyces lividans* xylanase 10A. Biochem J 2000, 350 Pt 3:933–941.

55. Kotake T, Dina S, Konishi T, Kaneko S, Igarashi K, Samejima M, Watanabe Y, Kimura K, Tsumuraya Y: Molecular cloning of a β-galactosidase from radish that specifically hydrolyzes β-(1->3)-and β-(1->6)-galactosyl residues of arabinogalactan protein. Plant Physiol 2005, 138:1563–1576.

56. Olson DG, Giannone RJ, Hettich RL, Lynd LR: Role of the *CipA* scaffoldin protein in cellulose solubilization, as determined by targeted gene deletion and complementation in *Clostridium thermocellum*. J Bacteriol 2013, 195:733–739.

57. Warnecke F, Luginbuhl P, Ivanova N, Ghassemian M, Richardson TH, Stege JT, Cayouette M, McHardy AC, Djordjevic G, Aboushadi N, et al: Metagenomic and functional analysis of hindgut microbiota of a wood-feeding higher termite. Nature 2007, 450:560–565.

58. Schwarz WH: The cellulosome and cellulose degradation by anaerobic bacteria. Appl Microbiol Biotechnol 2001, 56:634–649.

59. Kitago Y, Karita S, Watanabe N, Kamiya M, Aizawa T, Sakka K, Tanaka I: Crystal structure of Cel44A, a glycoside hydrolase family 44 endoglucanase from *Clostridium thermocellum*. J Biol Chem 2007, 282:35703–35711.

60. Blouzard J-C, Coutinho PM, Fierobe H-P, Henrissat B, Lignon S, Tardif C, Pagès S, de Philip P: Modulation of cellulosome composition in *Clostridium cellulolyticum*: Adaptation to the polysaccharide environment revealed by proteomic and carbohydrate-active enzyme analyses. Proteomics 2010, 10:541–554.

61. Yoshida S, Hespen CW, Beverly RL, Mackie RI, Cann IK: Domain analysis of a modular α-L-arabinofuranosidase with a unique carbohydrate binding strategy from the fiber-degrading bacterium *Fibrobacter succinogenes S85*. J Bacteriol 2010, 192:5424–5436.

62. Yoshida S, Mackie RI, Cann IK: Biochemical and domain analyses of *FSUAxe6B*, a modular acetyl xylan esterase, identify a unique carbohydrate binding module in *Fibrobacter succinogenes S85*. J Bacteriol 2010, 192:483–493.

63. Mackenzie AK, Pope PB, Pedersen HL, Gupta R, Morrison M, Willats WG, Eijsink VG: Two SusD-like proteins encoded within a polysaccharide utilization locus of an uncultured ruminant bacteroidetes phylotype bind strongly to cellulose. Appl Environ Microbiol 2012, 78:5935–5937.

64. Pope PB, Denman SE, Jones M, Tringe SG, Barry K, Malfatti SA, McHardy AC, Cheng JF, Hugenholtz P, McSweeney CS, Morrison M: Adaptation to herbivory by the Tammar wallaby includes bacterial and glycoside hydrolase profiles different from other herbivores. Proc Natl Acad Sci U S A 2010, 107:14793–14798.

65. Flint HJ, Bayer EA, Rincon MT, Lamed R, White BA: Polysaccharide utilization by gut bacteria: Potential for new insights from genomic analysis. Nat Rev Microbiol 2008, 6:121–131.

66. Naas AE, Mackenzie AK, J. M, Schückel J, Willats WGT, Eijsink VGH, Pope PB: Do Polysaccharide Utilization Loci represent an alternative mechanism for cellulose degradation? Under review (Submitted March 2014).

67. Morrison M, Pope PB, Denman SE, McSweeney CS: Plant biomass degradation by gut microbiomes: More of the same or something new? Curr Opin Biotechnol 2009, 20:358–363.

68. Martens EC, Lowe EC, Chiang H, Pudlo NA, Wu M, McNulty NP, Abbott DW, Henrissat B, Gilbert HJ, Bolam DN, Gordon JI: Recognition and degradation of plant cell wall polysaccharides by two human gut symbionts. PLoS Biol 2011, 9:e1001221.

69. Boraston AB, Bolam DN, Gilbert HJ, Davies GJ: Carbohydrate-binding modules: Fine-tuning polysaccharide recognition. Biochem J 2004, 382:769–781.

70. McNulty NP, Wu M, Erickson AR, Pan C, Erickson BK, Martens EC, Pudlo NA, Muegge BD, Henrissat B, Hettich RL, Gordon JI: Effects of diet on resource utilization by a model human gut microbiota containing Bacteroides cellulosilyticus WH2, a symbiont with an extensive glycobiome. PLoS Biol 2013, 11:e1001637.

71. Floudas D, Binder M, Riley R, Barry K, Blanchette RA, Henrissat B, Martinez AT, Otillar R, Spatafora JW, Yadav JS, et al: The paleozoic origin of enzymatic lignin decomposition reconstructed from 31 fungal genomes. Science 2012, 336:1715–1719.

72. Steyvers M, Griffiths T: Probabilistic topic models. In Handbook of latent semantic analysis. *Volume* 427. Edited by Landauer T, McNamara D, Dennis S, Kintsch W. Colorado, USA: Laurence Erlbaum; 2007: 427–440

73. Griffiths TL, Steyvers M: Finding scientific topics. Proc Natl Acad Sci U S A 2004, 101 Suppl 1:5228–5235.

74. Zhu W, Lomsadze A, Borodovsky M: Ab initio gene identification in metagenomic sequences. Nucleic Acids Res 2010, 38:e132.

75. McHardy AC, Martin HG, Tsirigos A, Hugenholtz P, Rigoutsos I: Accurate phylogenetic classification of variable-length DNA fragments. Nat Methods 2007, 4:63–72.

76. Patil KR, Haider P, Pope PB, Turnbaugh PJ, Morrison M, Scheffer T, McHardy AC: Taxonomic metagenome sequence assignment with structured output models. Nat Methods 2011, 8:191–192.

77. Yin Y, Mao X, Yang J, Chen X, Mao F, Xu Y: dbCAN: A web resource for automated carbohydrate-active enzyme annotation. Nucleic Acids Res 2012, 40:W445–451.

78. Eddy SR: Accelerated profile HMM searches. PLoS Comput Biol 2011, 7:e1002195.

79. Aravind L: Guilt by association: Contextual information in genome analysis. Genome Res 2000, 10:1074–1077.

80. Pagani I, Liolios K, Jansson J, Chen IM, Smirnova T, Nosrat B, Markowitz VM, Kyrpides NC: The genomes online database (GOLD) v.4: status of genomic and metagenomic projects and their associated metadata. Nucleic Acids Res 2012, 40:D571–579.

81. Levandowsky M, Winter D: Distance between sets. Nature 1971, 234:34–35.

82. Kuhn HW: The Hungarian method for the assignment problem. Nav Res Log 1955, 2:83–97.

83. Bron C, Kerbosch J: Algorithm 457: Finding all cliques of an undirected graph. Commun ACM 1973, 16:575–577.

84. Letunic I, Bork P: Interactive Tree Of Life v2: online annotation and display of phylogenetic trees made easy. Nucleic Acids Res 2011, 39:W475–W478.

85. Wilson DB: Three microbial strategies for plant cell wall degradation. Ann N Y Acad Sci 2008, 1125:289–297.

86. Suen G, Weimer PJ, Stevenson DM, Aylward FO, Boyum J, Deneke J, Drinkwater C, Ivanova NN, Mikhailova N, Chertkov O, et al: The complete genome sequence of *Fibrobacter succinogenes S85* reveals a cellulolytic and metabolic specialist. PLoS ONE 2011, 6:e18814.

87. McCartney L, Blake AW, Flint J, Bolam DN, Boraston AB, Gilbert HJ, Knox JP: Differential recognition of plant cell walls by microbial xylan-specific carbohydrate-binding modules. Proc Natl Acad Sci U S A 2006, 103:4765–4770.

